# Automating Academic Document Analysis with ChatGPT: A Mendeley Case

**DOI:** 10.1101/2024.03.18.585620

**Authors:** Mohamed Abuella

## Abstract

The management and organization of a large collection of academic documents is an important part of scientific research. This study explores the use of ChatGPT, a large language model from OpenAI, to extract insights from a large collection of academic documents stored in Mendeley. The study found that ChatGPT can be used to generate insightful graphs, concise summaries, and other tasks tailored to user needs. The study also demonstrated that ChatGPT can be used to analyze a large number of publications in PDF format. This suggests that ChatGPT could be a valuable tool for researchers who want to save time and effort by automating the analysis of their data. The GitHub repository for the source code and the output of this study is available at: https://github.com/MohamedAbuella/Analysis_Mendeley.

## I. Introduction

IN the realm of academic research, the diligent assembly of an extensive number of documents serves as a testament to the evolution of research knowledge. Over the years, since 2011, I have diligently built up a large collection of academic documents using Mendeley.

Multiple efforts have been made to derive more valuable information from this extensive archive.

The first attempt for analysis the Mendeley documents was during my doctoral study, when I encountered a review paper that employs text mining techniques to analyze 1000 publications in the field of solar energy forecasting [1]. Inspired by this approach, I considered applying similar techniques to analyze my Mendeley document collection. However, I faced challenges in implementing these techniques effectively.

Then, in the summer of 2022, I came across a paper by Wang et al. introduces the COVID-19 Open Research Dataset (CORD-19) [2]. The COVID-19 dataset comprises over 192,000 scholarly articles focusing on COVID-19, SARS-CoV-2, and related coronaviruses. The paper highlighted a collaborative effort among leading research groups, which, through Kaggle, launched a challenge and called upon artificial intelligence experts worldwide to contribute tools and approaches [3]. The aim was to assist the medical community in addressing high-priority scientific questions related to the ongoing health crisis.

After that, at the end of 2022 with the emergence of ChatGPT [4], a large language model capable of generating human-quality text and codes, I believe that was the opportune time to revisit my analysis of the collected Mendeley documents. ChatGPT’s ability to comprehend and process information from a wide range of code sources makes it an ideal tool with a query-based tool for obtaining coding assistance these documents into informative graphs and summaries.

In the course of this study, related works have been reviewed. Several studies have applied Natural Language Processing (NLP) for COVID-19 data analysis. For instance, Pratama et al. (2023) [5] conducted an exploratory data analysis of the COVID-19 Open Research Dataset (CORD-19) to identify trends and patterns in the data. Another study by Ekin et al. [6] clustered COVID-19 literature using NLP techniques to identify common themes and topics. Additionally, [7] introduced Llama 2, a family of open-access large language models for NLP tasks, which can be used for various COVID-19-related applications.

This study chronicles a personal project undertaken to harness the potential of ChatGPT as a utility for transforming Mendeley documents nto valuable and informative insights. The aim is to leverage ChatGPT’s capabilities to enhance the extraction and interpretation of meaningful information from the Mendeley document collection, thereby unlocking the full potential of this resource for research and knowledge development.

## II. Methodology

Flowchart depicting the systematic analysis process of Mendeley documents is shown in Figure 1.

**Fig. 1:**
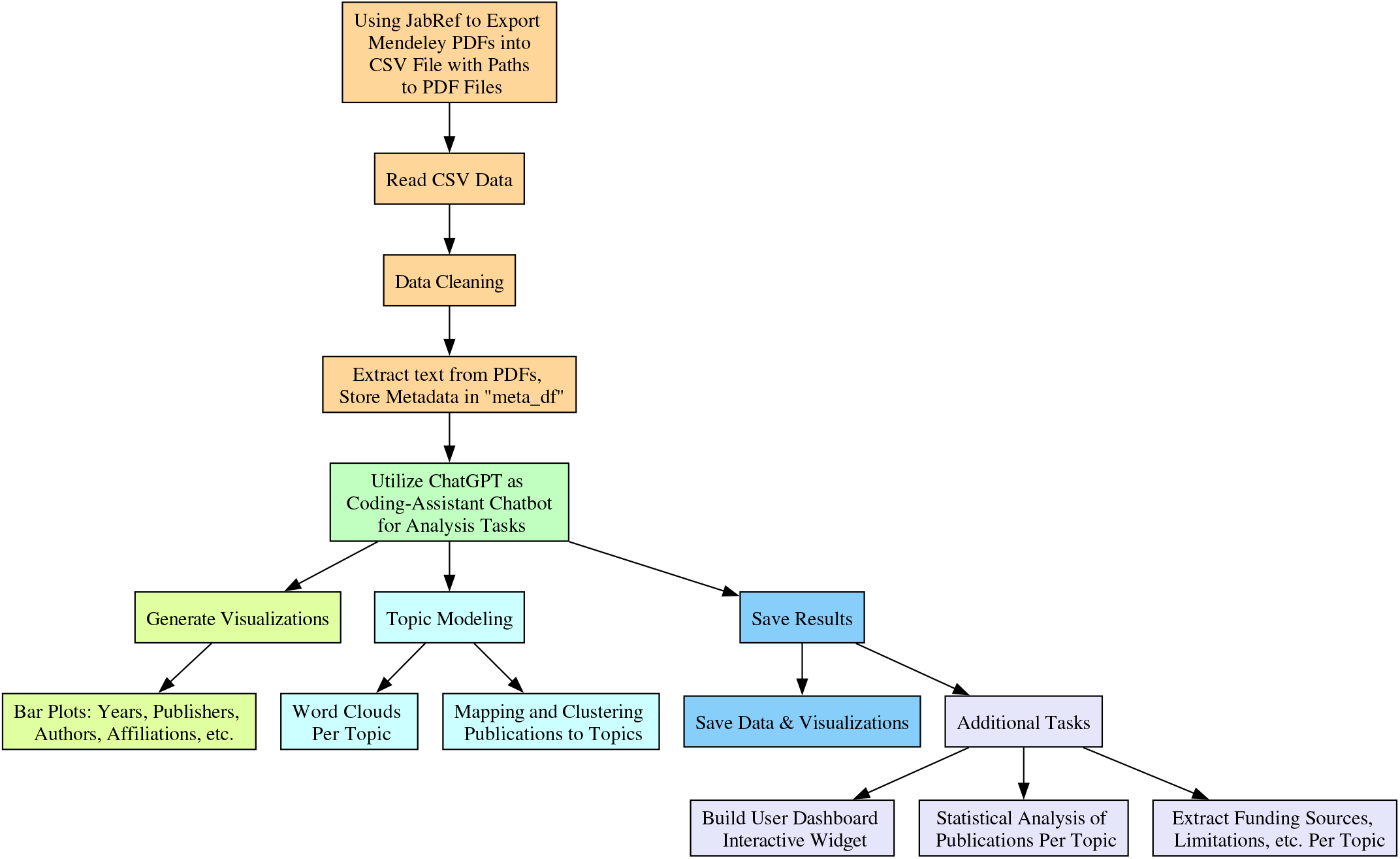
Flowchart for the proposed systematic analysis of Mendeley documents.

The initial step involves the extraction of data from Mendeley, comprising metadata such as titles, authors, and publication years. Subsequently, the contents of associated PDF files are processed using methods like **textract** and **PyPDF2** to capture valuable information embedded within the documents. Once the metadata are extracted from the documents, the pivotal role of the chatbot becomes apparent as it assists in coding and designing analysis tasks tailored to the user’s needs.

Keywords are extracted to facilitate a more granular analysis of the document content. The script not only extracts essential information but also performs topic modeling using NMF, enabling the identification of key themes within the document corpus.

## III. Results and discussion

The results of the analysis are presented through a series of visualizations, including bar plots depicting publication trends over the years, distribution of document types, and insights into prominent publishers and journals. Word clouds and interactive widgets offer an engaging visualization of user tags and enable detailed exploration of individual publications.

Furthermore, the incorporation of ChatGPT for topic modeling provides a comprehensive understanding of the over-arching themes within the document collection. The generated graphs and summaries offer a distilled representation of the knowledge landscape encapsulated in the Mendeley documents.

**Fig. 2:**
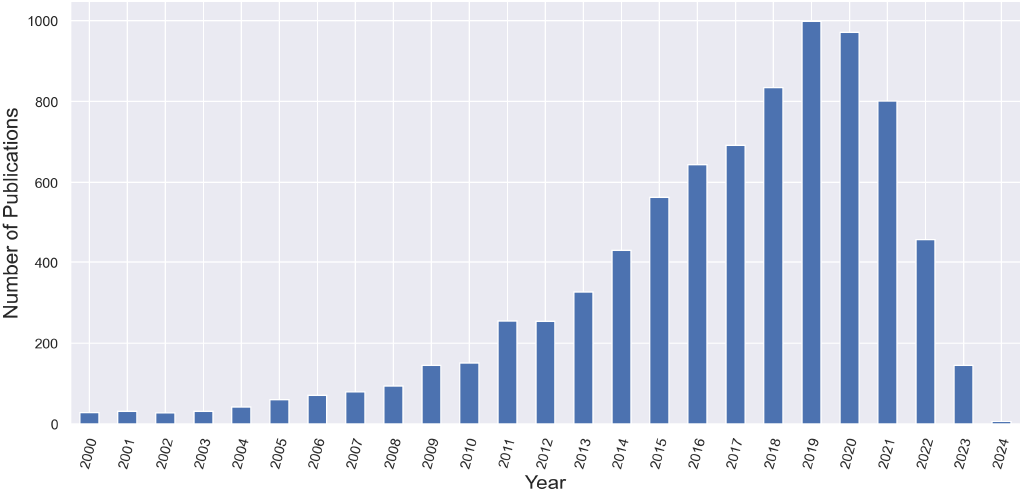
Barplot for number of publications per year.

**Fig. 3:**
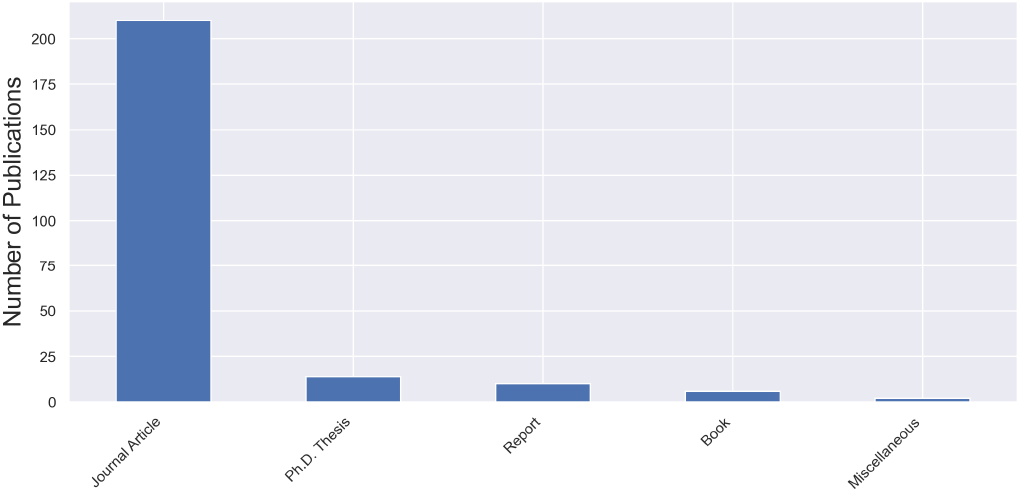
Barplot for number of publications per type.

**Fig. 4:**
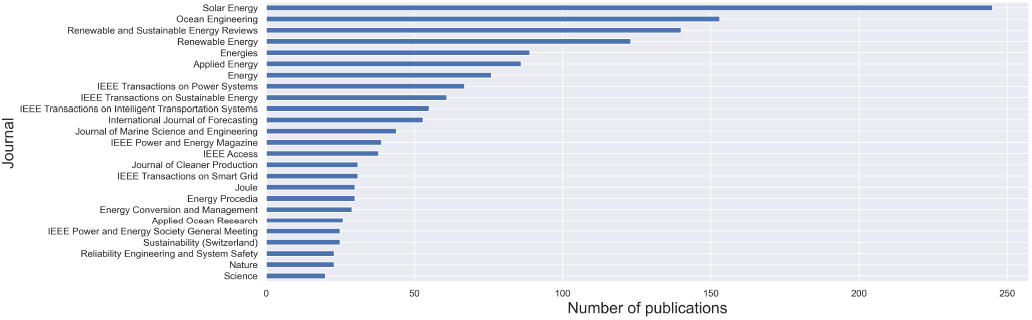
Barplot for number of publications per journal.

**Fig. 5:**
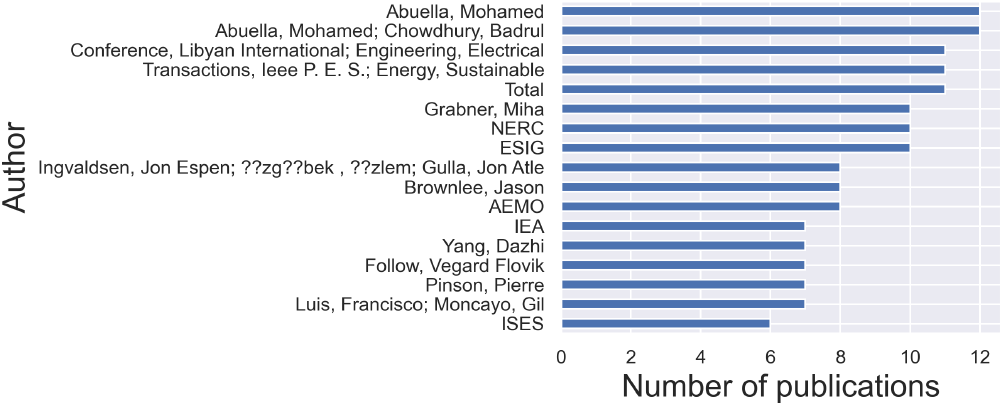
Barplot for number of publications per author.

**Fig. 6:**
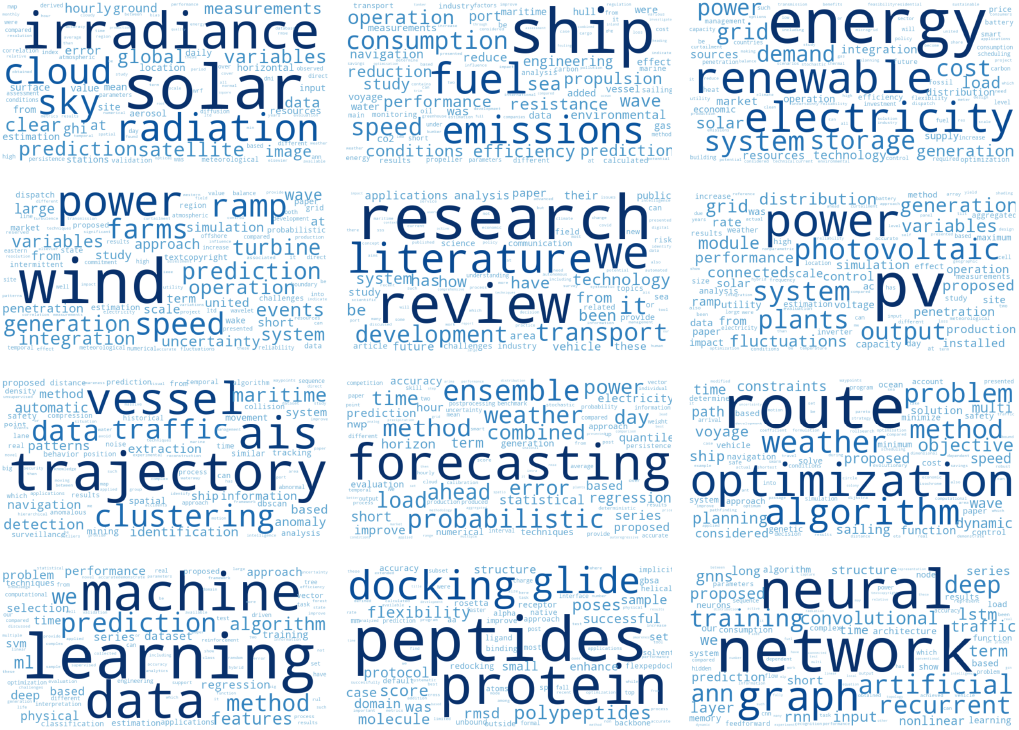
Word clouds of 12 topics in the publications.

**Fig. 7:**
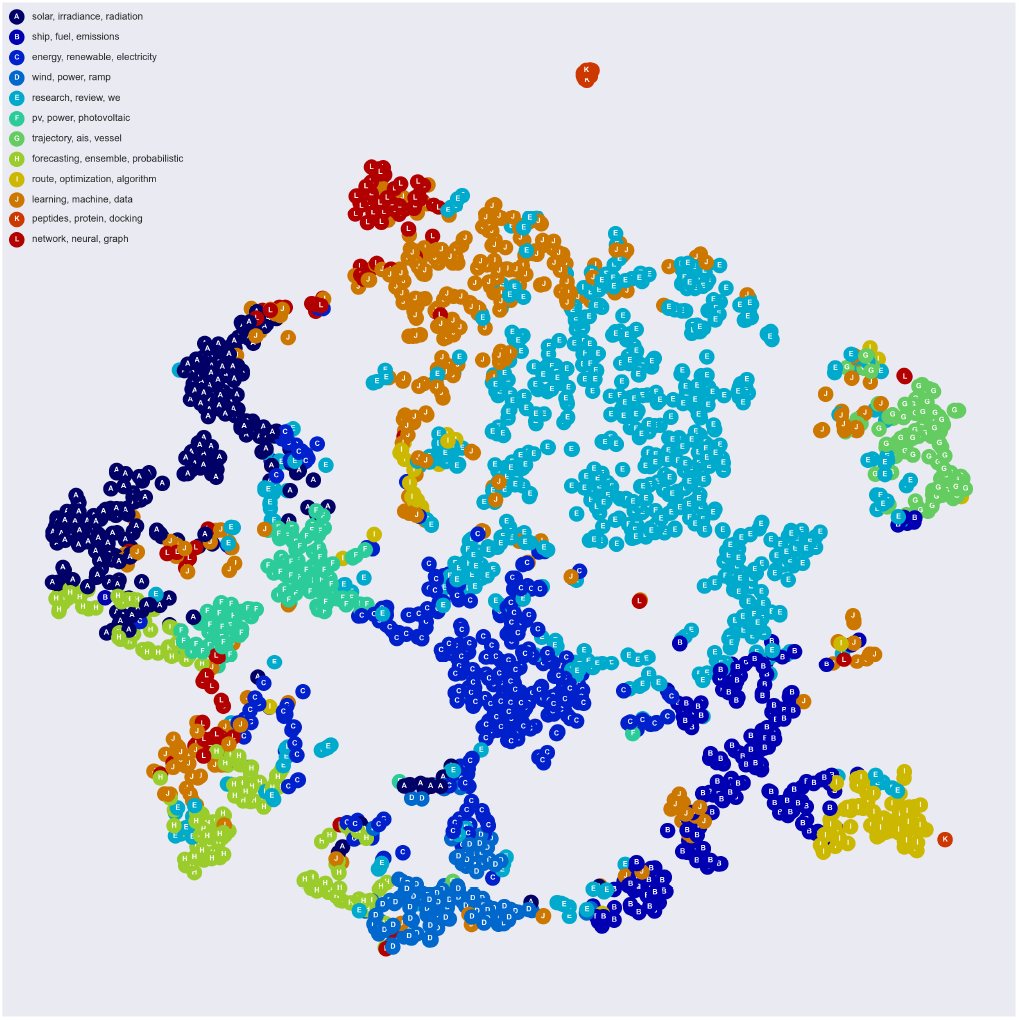
Map of the publications into the 12 topic clusters.

**Fig. 8:**
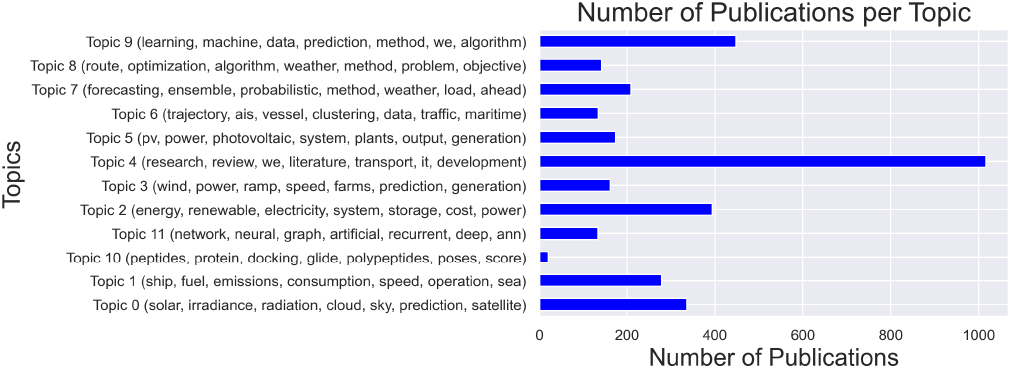
Number of publications per topic.

**Fig. 9:**
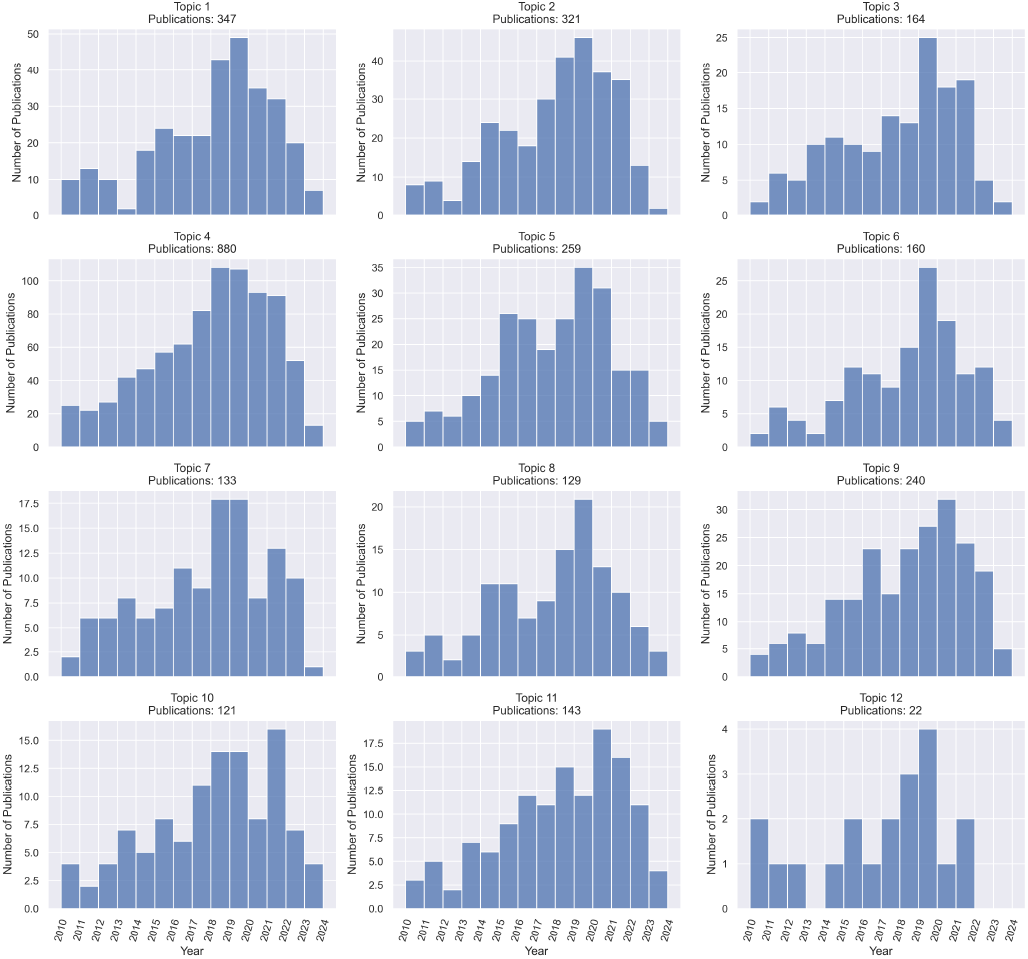
Number and year of publications per topic.

**Fig. 10:**
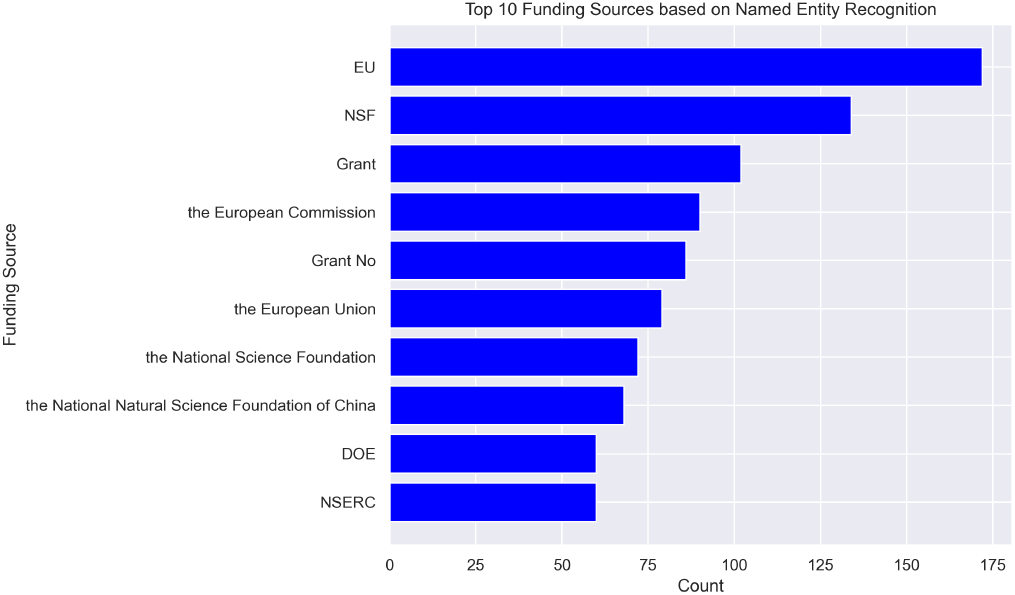
Top of funding sources for the publications.

## IV. Conclusion

This study showcases the successful application of ChatGPT as a coding utility for analyzing a large collection of Mendeley documents. The use of advanced natural language processing techniques, coupled with dynamic visualizations, transforms the traditional approach to document analysis. As a result, researchers and academics can derive valuable insights from their document repositories, enhancing the efficiency and depth of knowledge extraction.

In summary, the synergy between Mendeley and ChatGPT presents a compelling narrative of innovation and adaptability in the ever-evolving landscape of academic research. This project not only underscores the potential of AI-driven tools in document analysis but also serves as an inspiration for future endeavors aimed at unlocking the wealth of knowledge embedded in extensive document repositories.

Future work may involve further optimization of the search algorithm, exploration of additional natural language processing techniques, and extending the application to other domains. Moreover, integration of ChatGPT 4.0 plugins into the analysis pipeline could provide a dynamic and context-aware approach to understanding the nuances of the documents.

## Acknowledgment

The author wish to thank the diverse group at the Center for Applied Intelligent Systems Research (CAISR), Halmstad University, for helpful discussions.

## Appendix Supplementary Materials

The source codes that are implemented on Python 3.9.7 to produce the results are available at: https://github.com/MohamedAbuella/Analysis_Mendeley.

---

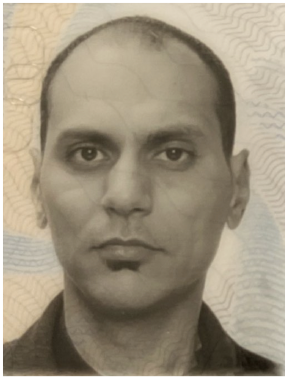

**Mohamed Abuella** received his M.S. and PhD degrees in Electrical and Computer Engineering from Southern Illinois University at Carbondale and University of North Carolina at Charlotte, in 2012 and 2018 respectively. He is a postdoctoral researcher at Halmstad University since 2022. His research interests include energy analytics and AI for sustainability.

